# mgikit: Demultiplexing toolkit for MGI fastq files

**DOI:** 10.1101/2024.01.09.574938

**Authors:** Ziad Al Bkhetan, Sen Wang

## Abstract

MGI sequencing is reported to be an inexpensive solution to obtain genomics information. There is a growing need for software and tools to analyse MGI’s outputs efficiently. *mgikit* is a tool collection to demultiplex MGI fastq data and reformat it effectively. The tool and its documentation are available at: https://sagc-bioinformatics.github.io/mgikit/.

## 1 Introduction

MGI Tech launched a series of new NGS equipment based on DNBSEQ technology. These sequencers have been reported to be similar or slightly less accurate than Illumina sequencers for different types of sequencing libraries. However, they are more cost-effective and have high throughput reaching ∼ 6 TB of data a day, as per the case of the T7 sequencer. These reasons pave the way for MGI sequencers to be widely utilised in the genomics field and therefore encourage the development of software that can analyse such data.

MGI sequencers output large fastq files with different read headers and file naming than Illumina outputs. The end of the reverse read (R2) for paired-end or the forward read (R1) for single-end is the read barcode that contains the sample’s indexes (i7 and i5) and unique molecule identifier (UMI). These indexes are used to demultiplex the data, that is, to assign the reads to the corresponding samples.

MGI tech has released the splitBarcode tool ^1^ to demultiplex MGI fastq. However, the tool does not recognise UMIs in the data, nor resolve the issue of different header and file naming formats that can be required by Illumina-based tools. In addition, it requires the user to know upfront where to find the indexes within the read barcode and it does not support multiple libraries within the same run smoothly.

In this application note, we provide *mgikit*, a software kit, to demultiplex MGI fastq data, detect barcode templates and generate demultiplexing and quality reports that can be converted to HTML reports through *mgikit-multiqc* plugin (available at https://github.com/sagc-bioinformatics/mgikit-multiqc) that integrates with the MultiQC tool [1]. *mgikit* is written in the RUST programming language. A comprehensive documentation and user guide are available at the tool web pages https://sagc-bioinformatics.github.io/mgikit/.

## 2 Results

### 2.1 fastq demultiplexing

For each read, the *mgikit* demultiplexer compares the read barcode with each sample index from the sample sheet and assigns the read to the sample with minimal mismatches. Reads that do not match with other samples are reported as undetermined. Reads that match multiple samples with equal errors are reported as ambiguous.

The ‘*demultiplex* ‘ command of *mgikit* is highly optimised to perform quick demultiplexing through buffering the output data to minimise writing-to-disk operations and dictionary-based data structure to achieve a quick and flexible search for matching barcodes. It requires one dictionary fetch for each index to find the associated sample and it accepts a user-defined threshold for allowed mismatches.

The main features provided by the *mgikit* demultiplexer compared to splitBarcode are highlighted in Table 1.

**Table 1:** Feature comparison between *mgikit* and splitBarcode.

| Feature | *mgikit* | splitBarcode |
| --- | --- | --- |
| Supports single-read, paired-end and 10X fastq input files | Yes | Yes |
| Supports mixed libraries | Yes | No |
| supports index diversity for samples | Yes | No |
| File naming and header format | Illumina and MGI format | MGI only |
| Support runs with UMI | Yes | No |
| demultiplexing and QC reports | text and html (through MultiQC plugin) | text |

### 2.2 Detection of barcode template

This utility is available using the ‘*template*’ command and it detects the barcode template for a data set. The barcode template represents the location of i7, i5 and UMI (if applicable) in the read barcode considering the reverse complementary forms. It tries all possible scenarios and selects the template that achieves the maximum number of matches.

### 2.3 Demultipexing reports

*mgikit* reports information on demultiplexing and read quality for each run. It also merges reports from multiple lanes into a comprehensive report for the whole run as well as for samples from different projects within the same run. *mgikit* reports can be parsed by the *mgikit-multiqc* plugin, as explained in the online documentation (https://github.com/sagc-bioinformatics/mgikit-multiqc), to generate user-friendly html reports integrated into multiqc [1] reports. *mgikit* provides ‘*report* ‘ functionality to merge multiple reports from multiple lanes together.

### 2.4 Performance indicators

The performance time of demultiplexing via *mgikit* (V0.1.2) and splitBarcode (V2.0.0) on the same datasets is reported in Table 2. Tests were done using a single thread on an Intel 2.80GHz machine with 32 GB RAM.

**Table 2:** Performance time (in minutes) evaluation and comparison on different datasets. DS01 and DS04 are 10 bp dual index, DS02 and DS3 is 8 bp dual index and DS05 is 8 bp single index. MK is *mgikit* while SP is splitBarcode. PE is paired-end while SE is single-end. In the case of single-end, the R2 file of the dataset is used alone for demultiplexing.

| Dataset | Reads | Samples | Length (bp) |  | Size (GB) |  | PE |  | SE |  |
| --- | --- | --- | --- | --- | --- | --- | --- | --- | --- | --- |
|  |  |  | R1 | R2 | R1 | R2 | MK | SB | MK | SB |
| DS01 | 298303014 | 102 | 300 | 320 | 76 | 85 | 71.5 | 84.6 | 37.2 | 45.5 |
| DS02 | 494667136 | 39 | 148 | 172 | 65 | 75 | 61.5 | 84 | 31.8 | 43.1 |
| DS03 | 506600595 | 29 | 100 | 124 | 46 | 55 | 43.5 | 65.5 | 30 | 35 |
| DS04 | 274567350 | 5 | 28 | 70 | 8.5 | 19 | 13 | 23.3 | 11.9 | 16.5 |
| DS05 | 500612381 | 64 | 50 | 8 | 22 | 5.5 | 12 | 31 | - | - |

## 3 Conclusion

We provide a tool kit for fastq demultiplexing and quality control reporting that can be utilised for the output of MGI sequencing machines and any other fastq datasets that have index information at the end of the read’s sequence. The tool can also be customised easily to suit different kinds of data. Further developments will focus on improving the performance time through CPU-parallel implementation as well as GPU-based functionality.

## Footnotes

1 https://github.com/MGI-tech-bioinformatics/splitBarcode

